# Reply to: “Model mimicry limits conclusions about neural tuning and can mistakenly imply unlikely priors”

**DOI:** 10.1101/2025.01.31.635589

**Authors:** Reuben Rideaux, Paul M Bays, William J Harrison

**Author notes:** Conflict of interest statement: The authors declare no competing interests.

## Abstract

In Harrison, Bays, and Rideaux (2023), we presented evidence from electroencephalographical recordings of humans that there is an over representation of horizontal orientations in the visual cortex. Wolf and Rademaker (2024) raise concerns about an analysis used in our study and provide an alternative explanation for our results. Here we address their concerns and provide additional magnetencephalography data supporting the conclusions of our original study.

## INTRODUCTION

A key goal of visual neuroscience is to understand how the physical properties of the world are represented by the brain. Efficient coding theory^1,2^ states that neural resources allocated to coding environmental features should be proportional to the frequency with which those features are found in nature. We recently found^3^ a horizontal bias in the neural representation of visual orientation, as measured in humans with electroencephalography (EEG). We then used *generative forward modelling*^4^, a method of comparing empirical neuroimaging recordings with matched simulated data produced by different population codes, to adjudicate between previously proposed and novel population codes of orientation in the visual cortex. Wolff and Rademaker^5^ replicated our main findings in their own data as well as in a re-analysis of our data: there is a horizontal bias in EEG measurements of orientation. They argue, however, that generative forward modelling has limited utility because it is susceptible to model mimicry, i.e. many different population codes could be responsible for the same pattern of EEG signals. Further, the authors propose an alternative explanation for the horizontal bias observed in EEG, involving an interaction between stimulus vignetting^6^ and a greater spatial representation of the horizontal meridian relative to the vertical meridian^7–9^. According to Wolff and Rademaker, this explanation is more plausible because it assumes equal representation of cardinal orientations and, in their view, there is little evidence supporting a horizontal bias in prior literature. Here we respond to these alternative explanations.

While we recognise that model mimicry presents a challenge in any inverse problem, we argue, contrary to Wolff and Rademaker, that rational constraints based on established neurophysiology can mitigate this risk. We will first clarify and expand on existing evidence that provides theoretical grounds for expecting a horizontal bias in neural representation, then explain why stimulus vignetting is unable to provide an alternative explanation for our results, and why Wolff and Rademaker’s findings for peripheral stimuli fail to challenge them. Finally, we highlight converging evidence for the horizontal bias obtained across multiple neuroimaging methods.

### Theoretical grounds and empirical evidence for a horizontal bias

There are two theoretical reasons to expect the existence of a horizontal bias in the human visual system, based on existing evidence. The first explanation was presented in our original study^3^: there are more horizontal orientations in natural visual scenes^10–13^. This statistical bias in natural visual environments is due in part to the prevalence of the horizon^14–17^, and as such is not confined to evolutionary history, although it is weaker in constructed environments. Our brains putatively mirror this natural anisotropy in the tuning properties of neurons in order to optimally process the environment^1,2^. The second theoretical explanation, which was not mentioned in our original study, relates to two well-known biases of the visual system: the radial orientation bias^18–21^ (an overrepresentation of neurons tuned to orientations that extend radially from fixation) and the horizontal meridian bias^7–9^ (a cortical overrepresentation of the visual field along the horizontal meridian). These biases were also discussed by Wolff and Rademaker, but they seem to have overlooked their implications. As shown in **Figure 1**, the interaction between these two biases produces an overrepresentation of horizontal orientation tuning in the visual cortex. This potential interaction between radial and spatial biases may reflect efficient coding of environmental statistics, but to the best of our knowledge we are the first to propose this hypothesis. Regardless, what is clear is that there are firm *a priori* grounds to expect a horizontal orientation bias in the human visual system.

**Figure 1.**
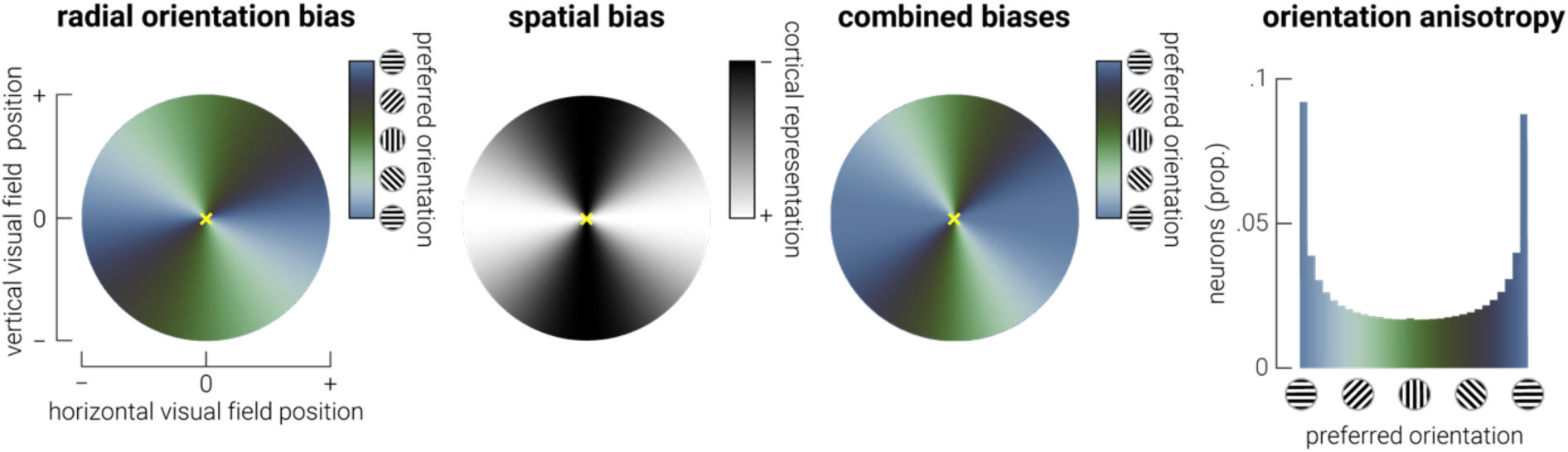
Interaction between radial orientation and spatial biases are sufficient to produce a horizontal orientation bias. Illustration demonstrating how well-established visual field biases could interact to produce a horizontal orientation bias. From left to right, the radial bias^18–21^ (overrepresentation of radially aligned orientations) and the spatial bias^7–9^ (overrepresentation of visual field along horizontal meridian) combine to produce an overrepresentation of horizontal orientation tuning preference in the visual cortex. That is, the spatial bias distorts the radial bias by expanding and contracting the regions around the horizontal and vertical meridians, respectively. The orientation anisotropy is evident in the radial plot (combined biases) as an increased representation of horizontal (blue colour) relative to the other orientations. Note, the yellow cross indicates the fovea.

Wolff and Rademaker (2024), in contrast, claim that there is little evidence for a horizontal bias in the existing literature. As discussed above, horizontal orientations are overrepresented in natural image statistics, which, under the efficient coding hypothesis, should be reflected in the encoding properties of the visual system. Indeed, in the psychophysics literature, this has been directly tested, and there has been consistent support for a horizontal bias for over three decades^14,22–25^. In our original article we highlighted evidence from several neurophysiological studies that points to a horizontal bias in the brain, including studies of mouse^26^, cat^27^, and non-human primate^18^. Other neurophysiological work also supports this bias^27–30^. However, we agree that that there are other neurophysiological studies, including those identified by Wolff and Rademaker, that have not observed a horizontal bias. Indeed, this lack of consensus in the literature was part of the motivation for our original study, and we hope it will inspire more direct examination of this important issue in future.

### Rational constraints mitigate the inverse problem

Wolff and Rademaker state that it is not possible to infer neural tuning from EEG without making assumptions about neurophysiology. We agree with this point. They also claim, however, that non-invasive imaging methods cannot give evidence about underlying neural causes^a^, and that incorporating established neurophysiology to constrain models “is not an option” because it would be relying on reverse inference^b^. These claims are overstated at best: much of our understanding of the encoding properties of the human visual cortex has been gained from indirect measures of latent neural processes (e.g., cross orientation suppression from fMRI measurements^31^). Indeed, Wolff and Rademaker themselves suggest that EEG can reveal anisotropic neural codes when they interpret results of their re-analysis as showing evidence of the oblique effect.

Generative forward modelling, like other analytic methods including computerized tomography (CT scans) and wavefield imaging (e.g., sonar), attempts to solve an inverse problem. In this respect, it is no different from other, more established, methods in that sensible constraints are required to reduce the range of possibilities and adjudicate between competing solutions. While in principle there are an infinite set of population codes (i.e. neural tuning functions) that could have produced the observed empirical results, in practice these population codes vary in their relative feasibility, as determined by physical laws (e.g., negative tuning functions aren’t possible) and existing empirical evidence (e.g., sharper tuning for obliques than cardinals directly contradicts neurophysiological evidence^30^). Moreover, we do not have to test all of them to decide between competing hypotheses, e.g. we can make simplifying assumptions including smoothly varying tuning preferences and widths, so long as smoothness is not the factor of interest. While we cannot assert that only a single population code produces the pattern of empirical results that were observed, we can use generative forward modelling to identify a tiny fraction of potential codes within the vast sea of possibilities, and then apply rational constraints to identify the most likely model.

Wolff and Rademaker show that in addition to a model with anisotropic tuning preferences, the empirical data can also be recapitulated by population codes with anisotropic tuning widths and gain (when the number of channels is also allowed to vary). The population code with anisotropic tuning width contradicts neurophysiological evidence^30^ and thus seems less plausible than the alternatives. As Wolff and Rademaker demonstrate, tuning preference and gain anisotropies produce highly correlated results. We agree with this limitation of our conclusions: in our population models, having two neurons tuned to the same feature produces the same population response as having a single neuron with twice the gain. For parsimony, in our study^3^ we limited the comparisons to those between tuning preference and width. However, while there is considerable empirical evidence of a cardinal/horizontal bias in the tuning preference of orientation selective neurons^18,26–30^, their response gain appears to be relatively isotropic^30^. It is notable that the population codes Wolff and Rademaker present as mimicking our results all also replicate our finding of an asymmetry between horizontal and vertical orientations, demonstrating that even when a wider parameter space is explored, some aspects of the code are necessary to explain the data.

### Differences between foveal and peripheral vision

It is encouraging that Wolff & Rademaker (2024) replicated our findings in their own foveal presentation data, and it is intriguing that they found no evidence of a horizontal bias in the periphery. There are many differences between the properties of the visual system that process information centrally and in the periphery, from the distribution of photoreceptors in the retina^32^ to the tuning properties of neurons in primary visual cortex, e.g., spatiotemporal frequency^33^ and binocular disparity^34^. Thus, given the known anisotropies that exist across the visual field, it seems reasonable to expect that biases for centrally presented features may be different than for those presented peripherally.

The results from stimuli presented to the left and right of fixation are particularly striking, given that the stimuli presented at these locations have considerable spatial overlap (∼20%) with those centered on fixation. Indeed, whether because of its interaction with a radial bias or stimulus vignetting, the over representation of the horizontal meridian should manifest in better decoding of horizontal orientations at the peripheral locations reported by Wolff and Rademaker. However, the discrepancy between the anisotropies observed at fixation and peripherally could be a result of other differences in experimental design. In particular, in Wolff and Rademaker’s experiment with peripheral presentation, two stimuli with different orientations were simultaneously presented to the left and right of fixation. It is unknown what effect presenting multiple concurrent stimuli might have on decoding of orientation from EEG recordings. We suggest that presence or absence of the horizontal bias in the periphery remains an open question, awaiting results of purpose-designed experiments.

### Stimulus vignetting unlikely to explain the horizontal bias

Wolff & Rademaker (2024) argue that the horizontal bias is due to an interaction between relatively reduced EEG signal-to-noise along the vertical midline, i.e., because of the aforementioned overrepresentation of visual field along the horizontal meridian, and stimulus vignetting^6^. The potential influence of stimulus vignetting has been raised in the context of multi-voxel pattern analysis of fMRI recordings, which is appropriate, as the spatial specificity of fMRI renders it particularly susceptible to this confound. That is, because activity produced by the stimulus and its border are represented separately by different voxels, a decoder can learn to classify the stimulus using information only from the border. By contrast, EEG recordings have low spatial resolution, so the activity at each sensor reflects the combined response to the stimulus and border. Thus, the influence of the border cannot be selectively increased, i.e., activity evoked by the border cannot be used selectively to decode orientation, but only as a component of the overall response to the stimulus. We therefore performed an image analysis to quantify the contribution of the border to the overall energy of a horizontally oriented target grating. This analysis is similar to the stimulus modelling^11,35^ used by Wolff & Rademaker (2024). Our conservative estimate shows that stimulus vignetting in our study produces a 3.9% increase in horizontal energy (**Fig. 2a**); however, when different spatial filters are used, this effect is either reduced or reversed such that vertical orientation energy dominates (see **Fig. S1**). Given the magnitude of the observed horizontal bias (horizontal/vertical decoding accuracy: 141%) is more than an order of magnitude larger than this, vignetting seems to be an unlikely explanation.

**Figure 2.**
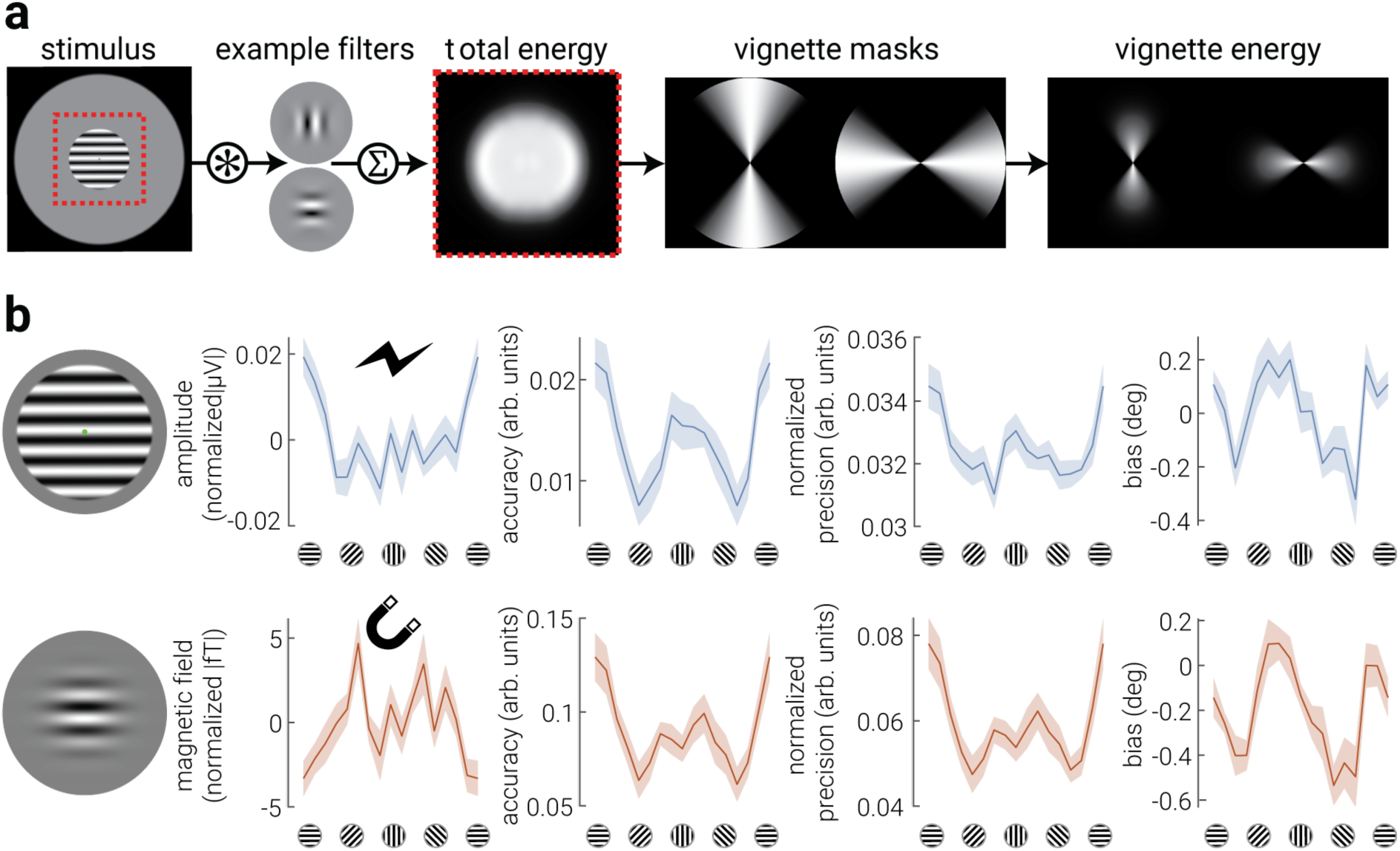
Stimulus vignetting is an unlikely explanation for the horizontal bias. **a**) Estimating the contribution of vignetting to the overall stimulus signal. We analysed the stimulus used in Harrison et al. (2023) and Myers et al. (2015) (stimulus shown in (**b**)) by convolving the stimuli with a bank of orientation and spatial frequency tuned filters, and then computing the extent of stimulus spread either parallel versus orthogonal to the stimulus orientation (see Supplementary Material for details description of analysis). The resulting difference between horizontal and vertical vignetting, as a percent of the total stimulus energy, was 3.9% and 0.5% for our study and Myers et al., respectively. **b**) From left to right: example stimuli, normalized absolute univariate response and corresponding decoding accuracy, precision, and bias, as a function of orientation, for Harrison et al. (2023) EEG dataset (top row, blue) and previously published Myers et al. (2015) MEG dataset (bottom row, orange). Shaded regions in (**b**) indicate ±SEM.

The relative energy contribution of stimulus vignetting increases with steeper border gradients^36^. Thus, if vignetting explains the horizontal bias, the bias we reported in our original study^3^ should be reduced or absent when a shallower border gradient is used. To test this, we re-analyzed a previously published dataset of magnetoencephalography (MEG) responses to centrally presented stimuli with considerably shallower borders^37^ (+0.5% horizontal energy from vignetting) using standard forward encoding (not generative forward modelling). In direct conflict with the vignetting account, we found a *larger* horizontal bias (**Fig. 2b**).

These findings provide compelling evidence against the vignetting explanation, but they also replicate the existence of the bias using a different neuroimaging modality. While MEG and EEG signals originate from the same neurophysiological processes, the biophysical properties of these signals are qualitatively different, e.g., one measures changes in electrical activity on the scalp while the other measures changes in the magnetic field. Indeed, in contrast to the results from EEG, we found that the univariate MEG responses were highest for obliques, and similar between horizontal and vertical orientations (**Fig. 2b**); as previous reported^38,39^. Despite this, we found the same pattern of decoding results, demonstrating a decoupling between the representational fidelity and the magnitude of the evoked response. These findings add to recent fMRI work^40^ that also show a strong horizontal bias, demonstrating converging evidence for a horizontal bias in human visual cortex across all major non-invasive neuroimaging modalities (EEG, MEG and fMRI).

### Concluding remarks

While vignetting was once thought to be a major confound for fMRI decoding^6^, recent work by the same group showed that orientation-specific BOLD activity does indeed reflect cortical tuning properties^41^. Similarly, our re-analysis of previously published data confirms that stimulus vignetting does not strongly influence orientation decoded from EEG/MEG. Rather, as we originally inferred^3^, the pattern of results we observed likely reflects a horizontal orientation bias in the visual cortex, and prior theoretical and empirical support for such a bias may have been overlooked. Further, in contrast to Wolff & Rademaker, we believe rejecting analytic methods simply because they attempt to solve an inverse problem is excessively conservative. We have shown that, as with other such methods, when rational constraints are applied, generative forward modelling can produce valuable insights into the population codes underlying neural responses.

## Acknowledgements

This work was supported by an Australian Research Council Discovery Early Career Researcher Award to RR (DE210100790). RR was also supported by a National Health and Medical Research Council (NHMRC; Australia) Investigator Grant (2026318).

## SUPPLEMENTARY MATERIALS

### Estimating the influence of stimulus vignetting

We analysed the potential contribution of vignetting to the overall stimulus energy using a standard filtering analysis (e.g. Harrison, 2021; Rideaux et al. 2021). An overview of the filtering is shown in **Figure 2a**. First, we filtered the target stimulus with a bank of 16 orientation and spatial frequency tuned quadrature pair filters. Orientations were linearly spaced between 0° and 168.75°. Changing the number of filters does not change the results. The spatial frequency was selected to match the period of the stimulus: 13 cycles per image for Harrison et al. (2023) and 16 cycles per image for Myers et al. (2015). Changing the spatial frequency tuning of the filter bank can invert the vignetting effect, a point to which we will return below. Each filter was convolved with the stimulus, and the resulting energies were summed across filter outputs. Filtering was performed in the frequency domain in MATLAB. We constructed two vignette masks: one that measures stimulus energy in the spatial region that extends parallel to the stimulus orientation, and one that measures stimulus energy in the spatial region that extends orthogonal to the stimulus orientation. Parallel and orthogonal vignette masks were raised cosine filters centred on 90° and 0°, respectively, with a bandwidth of 45°. Note that these filters are constructed in the spatial domain so that we could quantify spatial energy, unlike the oriented energy filters that were constructed in the frequency domain. We then found the dot product of the total stimulus energy and each vignette mask, thereby isolating energy that extends parallel or orthogonal to the stimulus orientation. Finally, the estimate of the contribution of vignetting to the total stimulus energy as a percent was calculated as:

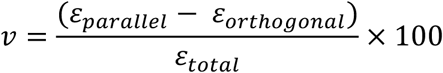

where *v* is the contribution of vignetting in percent, ε is stimulus energy from filters indicated by subscripts.

We estimated that vignetting contributes 3.9% to the overall signal based on the stimulus used in Harrison et al. (2023). Whereas Harrison et al. (2023) used a sinewave grating with a beveled edge, Myers et al. (2015) used a Gabor stimulus (4° diameter, 2 cycles/° spatial frequency) with a considerably smoother border profile (see **Figure 2b** for a visual comparison). As expected, the smoother border used by Myers et al. (2015) reduced its contribution to 0.5%. Changing the filter spatial frequency by ±1 octave changes the contribution of vignetting to the overall signal (see Supplementary **Figure S1**, below). For the targets used by Harrison et al. (2023), higher spatial frequency filters reduce the effect of vignetting, such that the difference in energy extending parallel to the stimulus orientation versus orthogonal to the stimulus contributes only 1% of the overall signal. Moreover, lower spatial frequency filters *reverse* the effect of vignetting, such that the energy extending orthogonal to the stimulus orientation is greater than energy extending parallel, contributing 27.4% of the overall signal when the filter is half the frequency as the target.

### Re-analysis of MEG dataset

To test whether vignetting could be driving the horizontal bias observed in Harrison et al. (2023), we re-analysed a previously published MEG dataset in which observers were presented with centrally positioned Gabor stimuli of varying orientation (Myers et al., 2015). The data were already pre-processed, as described in (Myers et al., 2015). Consistent with Harrison et al. (2023), we epoched the data from-50 to 450 ms around stimulus onset and included only posterior sensors in the analysis (EEG: occipital and parietal sensors; MEG: the 108 sensors between positions 1642 and 2543, according to the Elekta Neuromag electrodes scheme). For both EEG and MEG datasets, we sorted presentations into 16 evenly spaced orientations bins from 0° to 180° and calculated accuracy, precision, and bias in each bin, averaged across the epoch (see *Neural Decoding* section below for detailed description of analysis). In addition to these decoding metrics, we calculated the absolute univariate response as a function of orientation by averaging the absolute response (EEG: |µV|, MEG: |fT|) across the epoch. There were large individual differences in absolute univariate responses, so for the purpose of clarity, we normalized each participant’s average absolute univariate responses by subtracting the average.

### Neural Decoding

To characterise sensory representations of the stimuli, we used an inverted modelling approach to reconstruct stimulus orientation from the M/EEG recordings^31,42^. A theoretical (forward) model was nominated that described the measured activity in the M/EEG sensors given the orientation of the stimulus. The forward model was then used to obtain the inverse model that described the transformation from M/EEG sensor activity to stimulus orientation. The forward and inverse models were obtained using a ten-fold cross-validation approach in which 90% of the data were used to obtain the inverse model on which the remaining 10% were decoded.

Similar to previous work^43,44^, the forward model comprised five hypothetical channels, with evenly distributed idealized orientation preferences between 0° and 180°. Each channel consisted of a half-wave rectified sinusoid raised to the fifth power. The channels were arranged such that a tuning curve of any orientation preference could be expressed as a weighted sum of the five channels. The observed M/EEG activity for each presentation could be described by the following linear model:

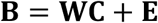

where **B** indicates the (*m* sensors × *n* presentations) M/EEG data, **W** is a weight matrix (*m* sensors × 5 channels) that describes the transformation from M/EEG activity to stimulus orientation, **C** denotes the hypothesized channel activities (5 channels × *n* presentations), and **E** indicates the residual errors.

To compute the inverse model, we estimated the weights that, when applied to the data, would reconstruct the underlying channel activities with the least error. In line with previous magnetencephalography work^45,46^, when computing the inverse model, we deviated from the forward model proposed by ^43^ by taking the noise covariance into account to optimize it for M/EEG data, given the high correlations between neighbouring sensors. We then estimated the weights that, when applied to the data, would reconstruct the underlying channel activities with the least error. Specifically, **B** and **C** were demeaned such that their average over presentations equalled zero for each sensor and channel, respectively. The inverse model was then estimated using either a subset selected through cross-fold validation) or all the data in one condition. The hypothetical responses of each of the five channels were calculated from the training data, resulting in the response row vector **c***_train,i_* of length *n_train_* presentations for each channel *i*. The weights on the sensors **w***_i_* were then obtained through least squares estimation for each channel:

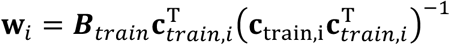

where ***B****_train_* indicates the (*m* sensors × *n_train_* presentations) training M/EEG data. Subsequently, the optimal spatial filter **v***_i_* to recover the activity of the *i*th channel was obtained as follows ^46^:

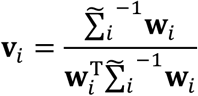

where 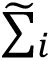 is the regularized covariance matrix for channel *i*. Incorporating the noise covariance in the filter estimation leads to the suppression of noise that arises from correlations between sensors. The noise covariance was estimated as follows:

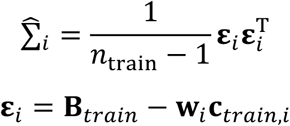

where *n_train_* is the number of training presentations. For optimal noise suppression, we improved this estimation by means of regularization by shrinkage using the analytically determined optimal shrinkage parameter^46^, yielding the regularized covariance matrix 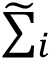.

For each presentation, we decoded orientation by converting the channel responses to polar form:

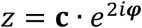

and calculating the estimated angle:

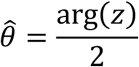

where **c** is a vector of channel responses and φ is the vector of angles at which the channels peak.

Stimulus orientation was sampled from continuous distributions, but to reliably characterize these features across their dimension, we grouped presentations into 16 evenly spaced bins from 0-180°. From the decoded orientation, we computed three estimates: *accuracy, precision*, and *bias*. Accuracy represented the similarity of the decoded orientation to the presented orientation^45^, and was expressed by projecting the mean resultant (averaged across presentations within the same stimulus orientation bin) of the difference between decoded and stimulus orientation onto a vector with 0°:

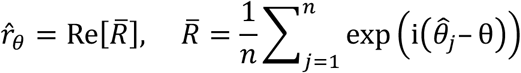

Precision was estimated by calculating the angular deviation^47^ of the decoded orientation within each orientation bin:

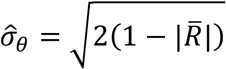

and normalized, such that values ranged from 0 to 1, where 0 indicates a uniform distribution of decoded orientation across all orientations (i.e., chance-level decoding) and 1 represents perfect consensus among decoded orientation:

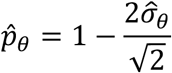

Bias was estimated by computing the circular mean of angular difference between the decoded and presented orientation:

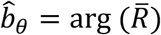

### Data availability statement

The EEG and MEG data re-analysed in this paper can be accessed at https://osf.io/5ba9y/ and https://doi.org/10.5061/dryad.m57sd, respectively.

### Code availability statement

The code used to perform the modelling can be accessed at https://osf.io/65dca/?view_only=88e12d759a67413ea4076a664a295798.

---

**Figure S1.**
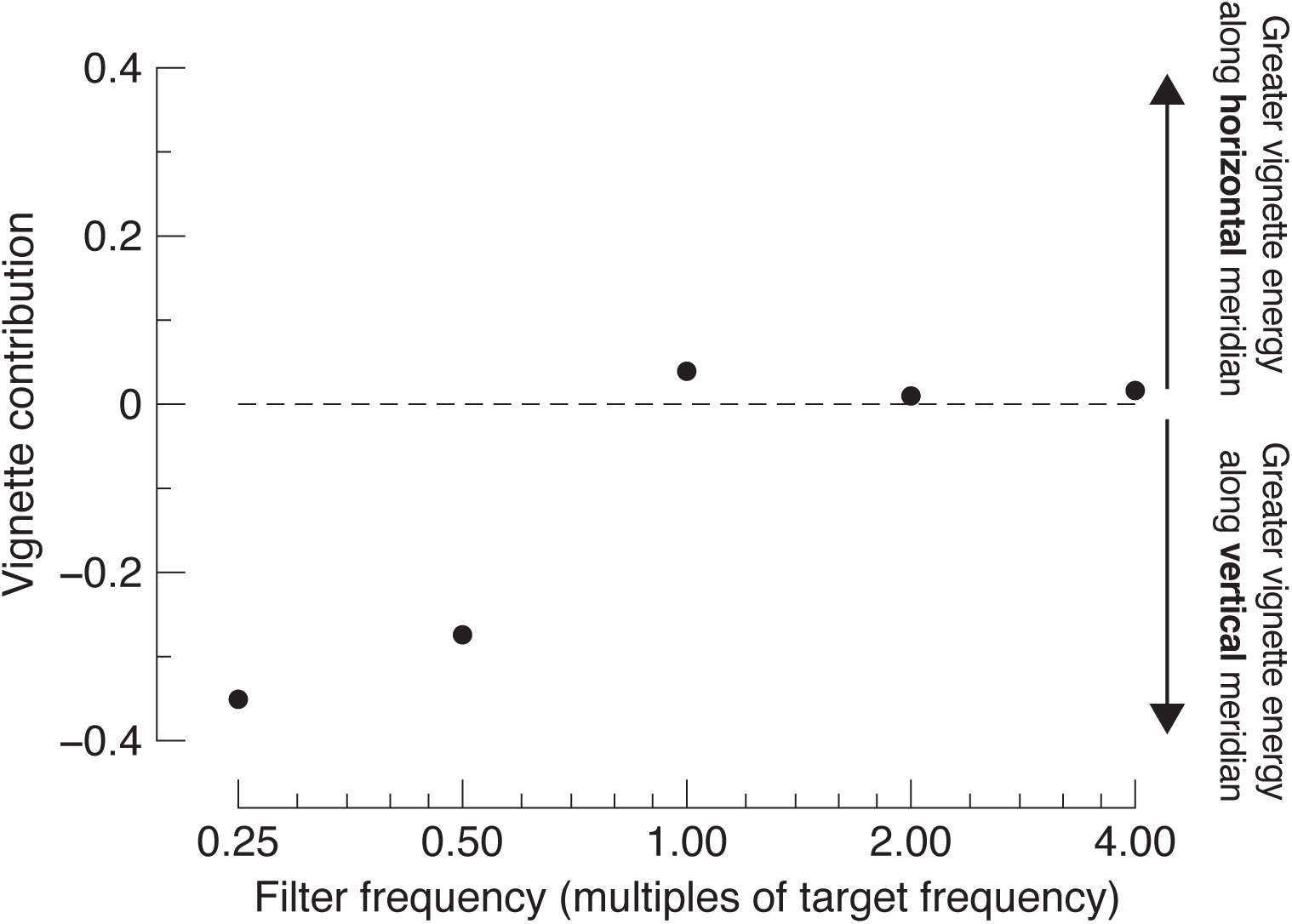
The contribution of the vignette energy to a horizontal grating, shown for filters of different spatial scales. Values above the dashed line indicate more horizontal vignetting than vertical, while values below the dashed line indicate more vertical vignetting than horizontal.

## Footnotes

a Specifically, they state, “The inverse problem is where an underlying cause cannot be inferred from a (measurable) e>ect, such as the inability to estimate neural causes from non-invasive imaging results.”

b Specifically, they state, “Excluding parts of this parameter space on the basis of previous physiology findings is not an option, as any subsequent claims about orientation tuning would amount to reverse inference.”

